# LP.8.1-directed COVID-19 mRNA vaccines durably boost neutralizing antibodies and mitigate ancestral immune imprinting

**DOI:** 10.64898/2026.01.03.697434

**Authors:** Ian A. Mellis, Madeline Wu, Hsiang Hong, Anthony Bowen, Kristin Daniel, Carmen Gherasim, Virginia M. Pierce, Michael T. Yin, Aubree Gordon, Yicheng Guo, David D. Ho

## Abstract

As SARS-CoV-2 evolves, it evades existing immunity elicited by exposure to earlier strains of the virus. In response, vaccine manufacturers have updated COVID-19 vaccines annually since 2022, though immune imprinting to the ancestral strain has blunted antibody responses to modern viral variants. In early 2025, the JN.1 subvariant LP.8.1 was dominant and manufacturers updated mRNA vaccine formulations to target LP.8.1 (LP.8.1 MV). However, by late 2025, other subvariants were dominant (XFG and NB.1.8.1) or emerging (e.g., PE.1.4, BA.3.2, PY.1.1.1) around the world. It is critical to understand the extent to which updated vaccine boosters elicit titers against both their target strain and recent variants. Further, it is important to quantify the extent to which immune imprinting continues to shape antiviral immune responses. Using pseudoviruses, we measured neutralizing antibody titers against a panel of 11 SARS-CoV-2 variants in serum samples from 36 adult participants in the United States before and approximately 1 month after LP.8.1 MV booster. We found that neutralizing antibody titers were substantially increased by the boost, with the greatest increases elicited against LP.8.1 and XFG. For the first time since 2022, post-boost titers were higher against the homologous vaccine target (LP.8.1) than against D614G (representing the ancestral strain). Combined, these results indicate that ancestral immune imprinting is mitigated to the greatest extent observed to date by LP.8.1 MV. Lastly, for a subset of participants, we measured neutralizing titers at approximately 4 months post-booster and found that LP.8.1-directed antibody titers were durable, with an estimated average half-life of approximately 66 days.

## Main Text

As SARS-CoV-2 evolves, it evades existing immunity elicited by exposure to earlier strains of the virus. In response, vaccine manufacturers have updated COVID-19 vaccines annually since 2022, though immune imprinting to the ancestral strain has blunted antibody responses to modern viral variants^1–3^. Throughout 2025, the global spread of SARS-CoV-2 has been dominated by a series of subvariants in the JN.1 sublineage, with intermittent concern for rarer divergent lineages, such as BA.3.2^4,5^. In early 2025, the JN.1 subvariant LP.8.1 was dominant in North America and other parts of the world. LP.8.1 is more resistant to serum antibody neutralization than early JN.1 subvariants, such as the KP.2 strain targeted by the 2024 mRNA vaccines, and it demonstrates greater affinity for the human ACE2 receptor^4^. In August 2025, the U.S. Food and Drug Administration authorized two updated mRNA vaccines (Pfizer-BioNTech and Moderna) based on the spike sequence of LP.8.1. In the United Kingdom, the European Union, and Japan, LP.8.1-based mRNA vaccines were also authorized. Here we report the acute boosting effect of updated LP.8.1 monovalent mRNA vaccines (LP.8.1 MV) on serum SARS-CoV-2-neutralizing antibodies in healthy adults in the United States. Others have reported peak boosting effects in other populations in Germany and Japan^6,7^. However, it remains essential to characterize boosting effects directly in the U.S. population, due to its different SARS-CoV-2 variant exposure, immunization, and COVID-19 incidence history^4,8,10^. Additionally, critically, because recent shifts in U.S. COVID-19 vaccination policy have created substantial uncertainty for both policymakers and frontline healthcare providers regarding age-based booster recommendations, we assess boosting effects in adults across multiple age ranges, including those younger than 50 years-old, 50 to 64 years-old, and 65 years or older. Lastly, it is important to characterize the durability of LP.8.1 MV boosting effects, which has not yet been discussed by others in the literature.

Since the authorization of the updated LP.8.1 MV boosters, SARS-CoV-2 has evolved beyond LP.8.1, with JN.1 subvariants XFG and NB.1.8.1 becoming dominant globally. Other JN.1 subvariants, such as PY.1.1.1 in North America, PE.1.4 in Oceania, as well as the highly divergent BA.3-descendant BA.3.2 sublineage, mostly in Australia and Germany, are also growing in incidence (**Figure S1**). XFG, NB.1.8.1, PE.1.4, and PY.1.1.1 contain 8, 5, 10, and 10 spike amino-acid differences relative to LP.8.1, respectively, and BA.3.2 has dozens of amino-acid differences, in a separate viral sublineage entirely (**Figure 1A**). It is critical to assess the effectiveness of the updated LP.8.1 MV boosters on neutralizing antibodies in human sera against recently dominant subvariants and those that are of growing concern. Furthermore, it is important to understand the extent to which ancestral immune imprinting still shapes humoral immunity against circulating SARS-CoV-2 variants.

**Figure 1.**
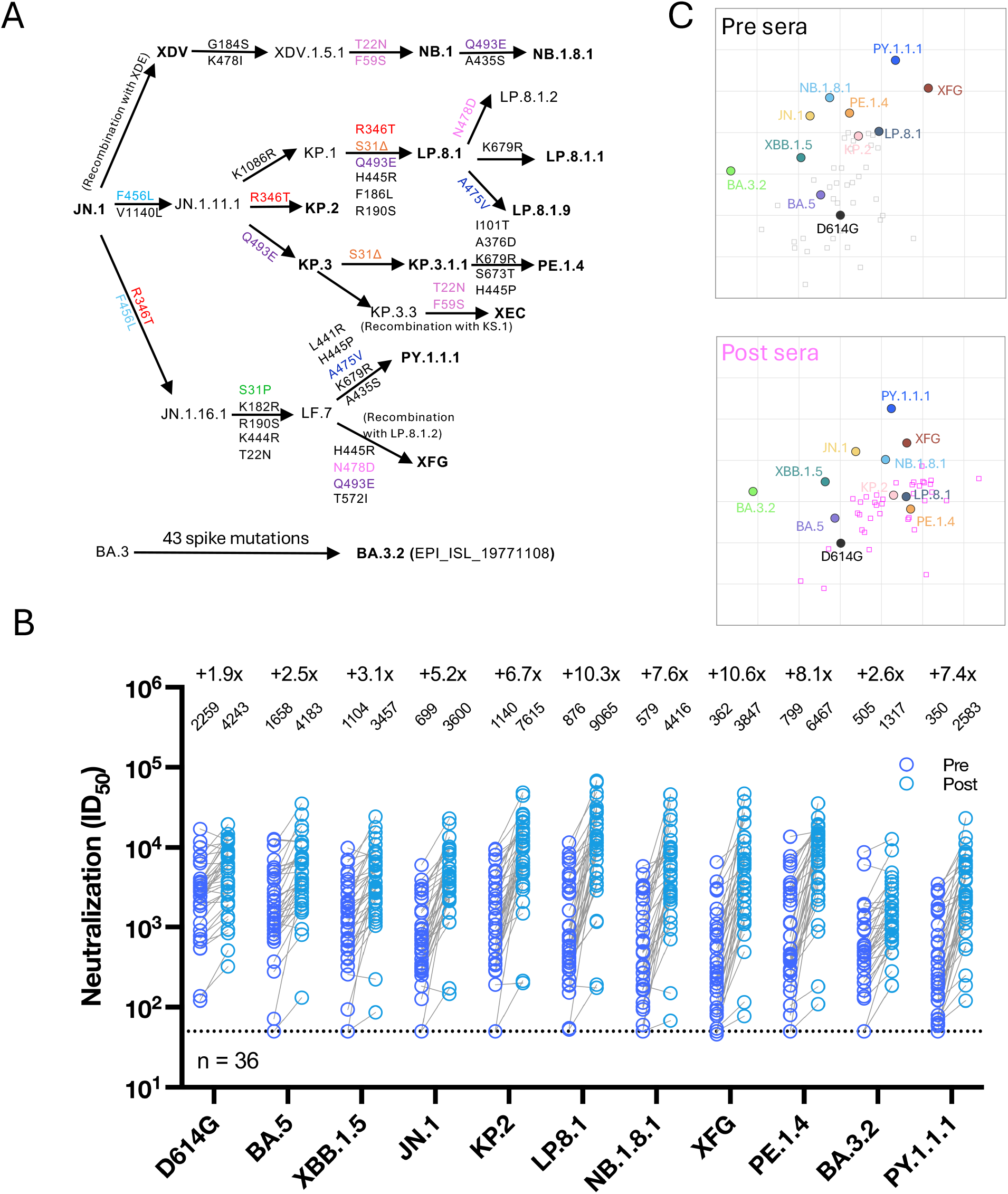
SARS-CoV-2-neutralizing antibody titers before and after a LP.8.1 monovalent mRNA vaccine booster. **A**. Spike mutations of the indicated JN.1 sublineages. Δ, deletion. **B**. Serum virus-neutralizing titers (ID_50_) against the indicated SARS-CoV-2 pseudoviruses and the fold changes between pre- and post-booster serum samples are shown. Geometric mean titers (GMT) are shown above each sample set, and the fold change from pre- to post-booster is shown above GMTs. MV, monovalent vaccine. n, sample size. The dotted line represents the assay limit of detection (LOD) of 50. C. Antigenic mapping of SARS-CoV-2 variants before and after a LP.8.1 monovalent mRNA vaccine booster.

To assess serum neutralizing antibodies elicited by LP.8.1 MV boosters, we generated vesicular stomatitis virus (VSV)-based pseudoviruses for historic and circulating SARS-CoV-2 variants: D614G, BA.5, XBB.1.5, JN.1, KP.2, LP.8.1, NB.1.8.1, XFG, PE.1.4, BA.3.2, and PY.1.1.1. We then performed pseudovirus neutralization assays using serum samples collected before and approximately 1 month (mean 28.6 days, range 21-45 days) after vaccination (**Tables S1, S2**). Vaccination boosted serum virus-neutralizing 50% inhibitory dilution (ID_50_) titers by 1.9-fold against D614G, 2.5-fold against BA.5, and 3.1-fold against XBB.1.5 (**Figure 1B**), reflecting persistent but modest back-boosting comparable to last year’s KP.2 booster. However, LP.8.1 MV induced a more substantial boost (5.2- to 10.6-fold) against subvariants in the JN.1 lineage: JN.1, KP.2, LP.8.1, NB.1.8.1, XFG, PE.1.4, and PY.1.1.1, with the greatest increases against LP.8.1 (10.3-fold) and the present dominant strain XFG (10.6-fold). Pre-boost titers (Geometric Mean ID_50_ [GMT]), reflective of much of the U.S. population, varied widely across variants (350 to 2,259), with highest titers against D614G and lowest against PY.1.1.1. These pre-boost titers suggest that a substantial fraction of the population may have low baseline protection from infection by variants such as BA.3.2 (GMT = 505) and XFG (GMT = 362), based on our prior studies of neutralizing antibody correlates of protection from earlier Omicron variants in other cohorts^9^. However, post-boost virus-neutralizing titers were quite robust (1,317 to 9,065), with the highest value observed for LP.8.1 (GMT = 9,065), correlated with greater protection in earlier studies. LP.8.1 MV also induced a modest boost (2.6-fold) against the highly divergent BA.3.2 strain (post-boost GMT = 1,317). This marks the first time that we have observed post-boost titers higher against the homologous target strain than against the ancestral D614G. In previous years, BA.5 bivalent (BA.5 BV), XBB.1.5 monovalent (XBB.1.5 MV), and KP.2 monovalent (KP.2 MV) mRNA vaccines all elicited post-boost titers that were highest against D614G^1–3^.

To assess antigenicity of the tested variants, we performed antigenic mapping using the pre-boost and post-boost sample sets separately (**Figure 1C**). Antigenic distances between post-boost sera and viral variants were reduced compared with pre-vaccination sera, confirming an overall boosting effect of the LP.8.1 MV. In addition, the relative positions of variants in the antigenic map were largely preserved before and after boosting, suggesting that neutralization correlation patterns across variants did not substantially change. However, serum samples in the pre-boost set were embedded primarily near D614G and BA.5 whereas in the post-boost set they shifted toward LP.8.1, concordant with maximum post-boost titers against LP.8.1. Combined with dramatic homologous LP.8.1-boosting and low back-boosting to D614G, these results suggest that LP.8.1 MV vaccination substantially mitigated ancestral immune imprinting, possibly to the greatest extent observed for any updated vaccine to date. This mitigation of imprinting is associated both with increased genetic distance of LP.8.1 from the imprinted ancestral strain and with an accumulated larger number of monovalent Omicron subvariant exposures in this cohort compared with earlier studies. For example, in this cohort the average total number of monovalent Omicron-directed vaccine doses (i.e., XBB.1.5 MV, KP.2 MV, and LP.8.1 MV combined) was 2.6, while in our prior study of KP.2 MV boosters the average number was 1.8^3^.

To quantify the effects of immune imprinting on neutralizing antibody responses elicited by different boosters, we compared LP.8.1 MV booster-elicited titers to titers we previously measured in earlier cohorts boosted with BA.5 BV, XBB.1.5 MV, or KP.2 MV [1-3]. First, we compared the fold-changes in titers against the imprinted D614G strain and the titers against the target strain of each vaccine (i.e., BA.5 for BA.5 BV, XBB.1.5 for XBB.1.5 MV, and KP.2 for KP.2 MV). Consistent with prior results, we found that monovalent Omicron-subvariant-directed vaccines elicited larger target fold changes (7.8X-13X) than ancestral-directed titer back-boosts (1.6-3.2X), particularly when compared to the BA.5 BV formulation (2.6X target-directed, 1.9X ancestral-directed; Figure 2A). Next, we compared post-boost titers against the respective target strains of each of the four boosters versus post-boost titers against D614G. We found substantially larger fold differences in post-boost titers after LP.8.1 MV (2.1X, LP.8.1 vs. D614G) compared to earlier boosters (0.09X-0.22X, target vs. D614G), suggesting that the strength of LP.8.1 MV booster-elicited target-directed responses is not limited by immune imprinting as much as in earlier years (Figure 2B).

**Figure 2.**
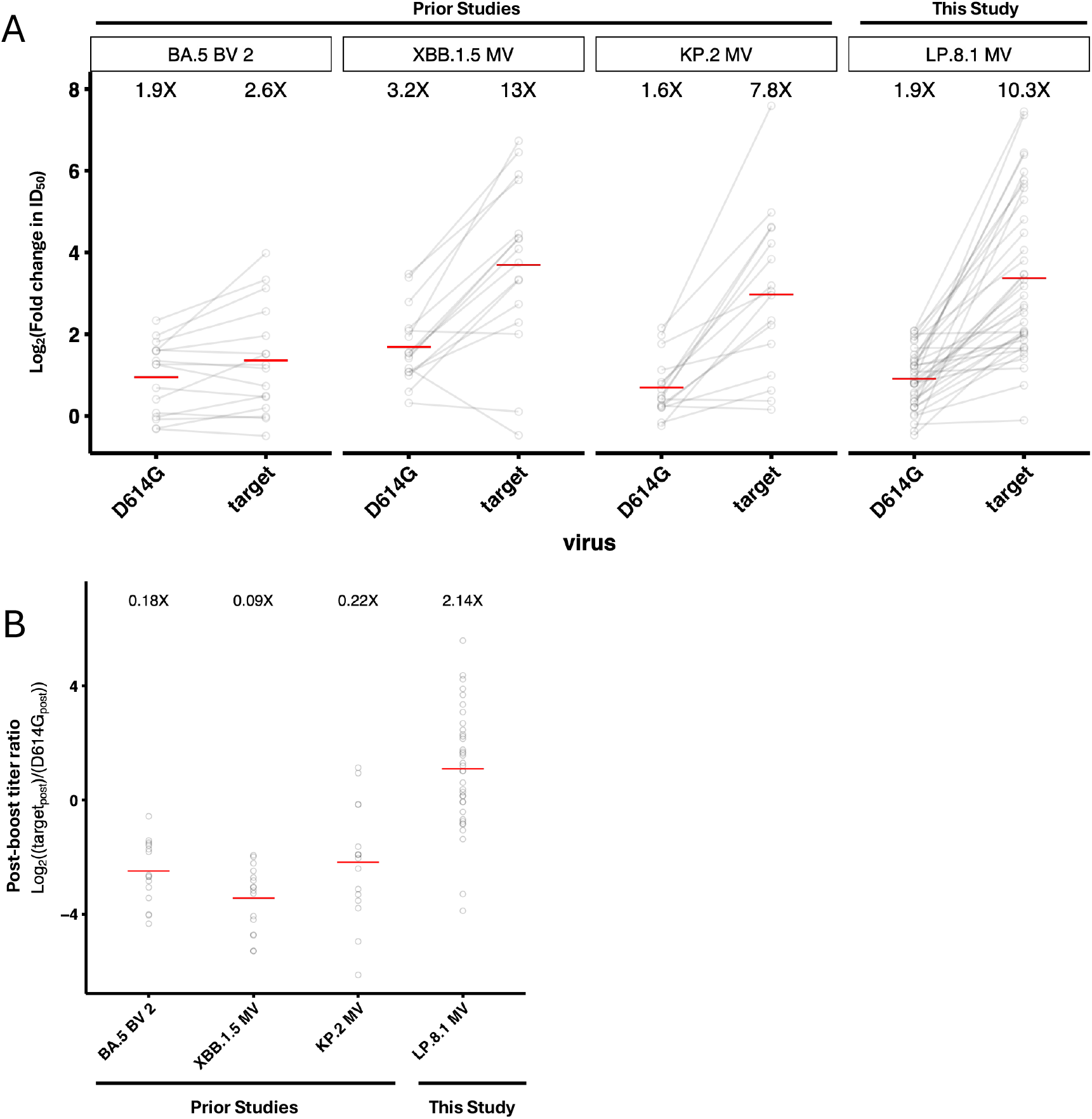
Mitigation of immune imprinting by Omicron-subvariant-directed monovalent mRNA vaccines. A. Fold change in ID50 against D614G or the target variant of each vaccine, as measured in pseudovirus neutralization assays. BA.5 BV 2, second bivalent wildtype/BA.5 bivalent booster; XBB.1.5 MV, XBB.1.5 monovalent vaccine booster; and KP.2 MV, KP.2 monovalent vaccine booster data from previously published studies (citations). Red lines indicate geometric mean fold change in titer, which is noted above each group of results. **B**. Fold differences in post-boost ID50 against the target variant of each vaccine vs. against D614G, using the same data sources (target_post_ / D614G_post_). Red lines indicate geometric mean fold difference, which is also noted above each group of results.

To better understand how LP.8.1 MV boosting performs across the adult age spectrum under current age-focused vaccination policies, we stratified participants into three age groups. The 36 adult participants in this study spanned multiple age ranges: 18 participants were between 18-49 years-old, 10 were 50-64 years-old, and 8 were 65 years-old or older (full cohort mean 47.3, range 19 to 80). All age range groups demonstrated similar trends in boosting: the fold increases in GMT against D614G were 1.9X, 1.9X, and 1.8X, respectively, and fold increases against LP.8.1 were 9.7X, 8.9X, and 14.4X, respectively (**Figure S2**). Post-boost titers were potent and broad cohort-wide, but titers against all JN.1 subvariants were highest overall in the oldest age group. Post-boost GMTs against D614G were 4,810, 3,947, and 3,504, respectively, while those against LP.8.1 were 6,795, 7,551, and 21,787, respectively. These observed levels of neutralizing antibodies were predictive of protection against symptomatic COVID-19 in prior clinical studies, but the precise titers that correlate with protection will vary according to the neutralization assays employed. Importantly, in addition to different numbers of participants in each age group, there were differences in numbers of reported lifetime COVID-19 vaccine doses: 5.9, 7, or 8 total doses respectively, of which an average of 3, 3.4, or 3.6 were of the wild-type formulation, 0.6, 1, or 0.9 were wild-type/BA.5 bivalent, and 2.3, 2.6, or 3.5 doses of monovalent Omicron-subvariant-directed vaccines **(Table S1)**. Therefore, in the tested cohort, post-boost titer differences between age groups may be confounded with vaccination history differences.

Lastly, to assess the durability of antibody responses to LP.8.1 MV boosters, we measured neutralizing titers against a panel of 8 SARS-CoV-2 variants at approximately 1 and 4 months post-boost for 11 participants. We found that titers were durable over the study period in this small cohort, with half lives per variant ranging from >43 days against XFG and >66 days against LP.8.1 to >292 days against D614G (Figure 3). It is intriguing to note potential differences in variant-specific titer decay rates, with cross-reactive imprinting-associated titers, e.g., for D614G and perhaps also cross-reactive to BA.3.2, showing slower decay than newly elicited target-variant-specific anti-LP.8.1 and -XFG titers.

**Figure 3.**
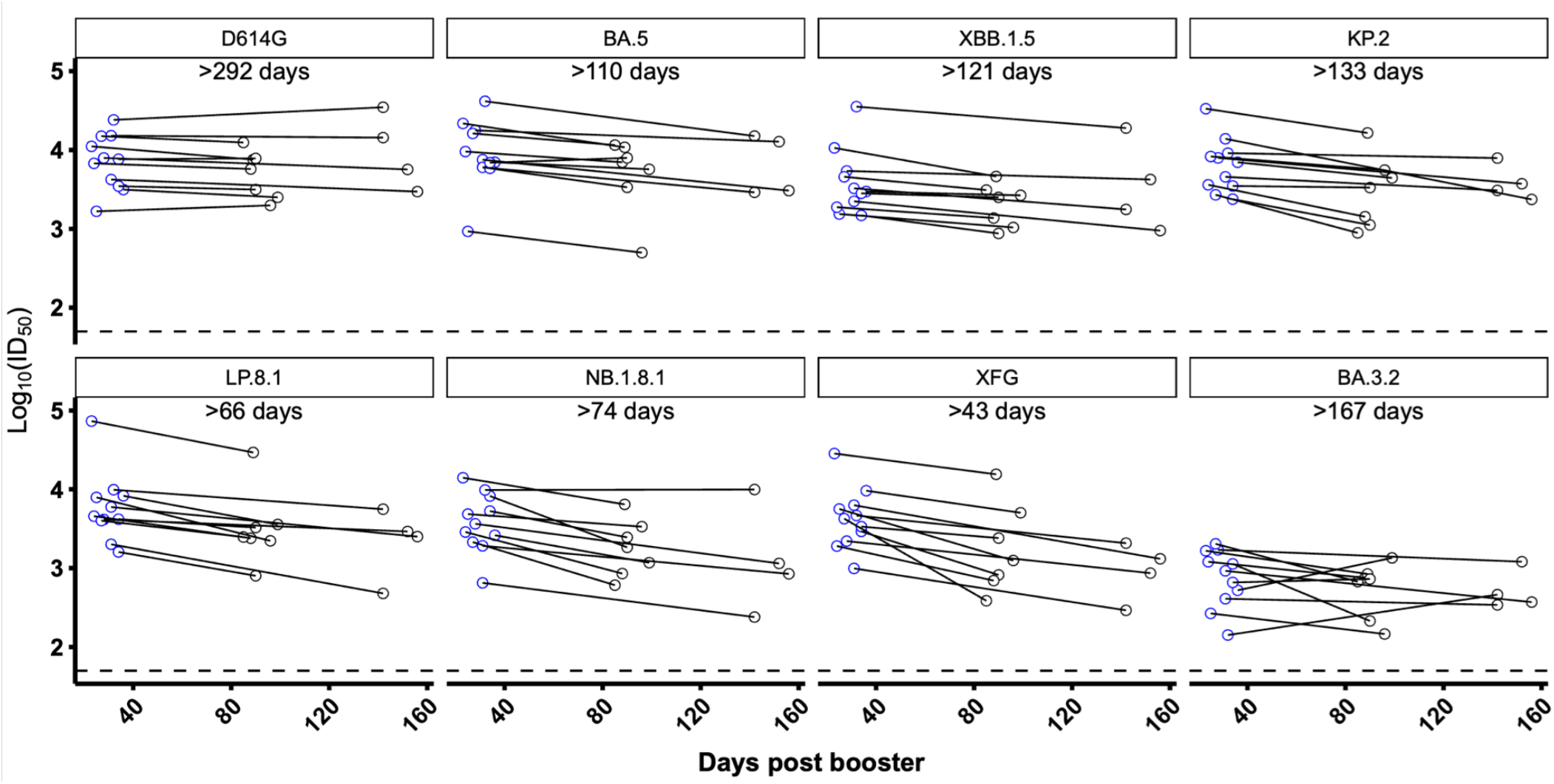
Durability of neutralizing antibody titers after LP.8.1 MV booster. Serum virus-neutralizing titers (ID_50_) for 11 participants against the 8 indicated SARS-CoV-2 pseudoviruses. Estimated half-life shown for each variant-directed titer. The dotted line represents the assay limit of detection (LOD) of 50.

In summary, a LP.8.1 MV booster elicits a robust and durable neutralizing antibody response against LP.8.1 and other JN.1 subvariants in healthy U.S. adults across age ranges. In this cohort, immune imprinting to the ancestral strain of SARS-CoV-2 was noticeably mitigated irrespective of age. Notably, and in contrast to prior reports in populations with different prior exposure histories, we observe highest post-boost titers against the homologous strain, LP.8.1. Overall, we observe similarly broad and potent booster-elicited neutralizing antibody responses. Furthermore, we show for the first time that LP.8.1 boosts titers against other emerging variants of concern, including PE.1.4 and PY.1.1.1, and we characterize durability of LP.8.1 MV-elicited neutralizing antibody titers for the first time. The durability of titers against circulating and historical variants should continue to be monitored over the coming months, as should the clinical effectiveness of the vaccines in protecting from symptomatic infections and severe disease. Together, these results highlight how updating COVID-19 vaccines to combat contemporary viral variants is a promising and important strategy for mitigating suboptimal ancestral-strain-focused immune responses elicited by earlier exposures.

