## Supplemental Appendix for "LP.8.1-directed COVID-19 mRNA vaccines durably boost neutralizing antibodies and mitigate ancestral immune imprinting"

### Supplemental Methods

#### *Clinical cohorts*

Serum samples were collected as part of the VIVA study at the University of Michigan and as part of the "COVID-19 Persistence and Immunology Cohort (C-PIC)" study at Columbia University. Specimens were obtained following participant informed consent and in adherence to the protocols approved by the IRBs of University of Michigan Medical School (protocol HUM00232359) and Columbia University (protocol AAAS9722).

In this study, serum samples were collected from individuals who had been administered the LP.8.1 monovalent vaccine booster (LP.8.1 MV). Serum samples were collected both before and after booster administration. The majority of the study subjects were female (77.8%) with an average age of 47.3 years. Serum samples were collected, on average, 6.1 days pre LP.8.1 MV booster and 28.6 days post LP.8.1 MV booster. Pre-, 1-month post-, and 4-month post-boost samples were examined by anti-nucleoprotein (NP) ELISA to check for unreported intervening SARS-CoV-2 infections. Further demographic details, vaccination status, and serum collected timelines can be found summarized in **Tables S1, S2, S3, and S4**.

#### *Cell lines*

HEK293T (ATCC, CRL-3216) cells and Vero-E6 cells (ATCC, CRL-1586) were cultured in complete culture medium: Dulbecco's Modified Eagle's Medium (DMEM) that contained 10% heat-inactivated fetal bovine serum (FBS) and 1% penicillin-streptomycin (PS). All cell lines were cultured in an atmosphere of 5% $CO_2$ at 37°C.

#### *SARS-CoV-2 spike plasmids*

The spike constructs of D614G, BA.5, XBB.1.5, JN.1, KP.2, LP.8.1, XFG, and NB.1.8.1 were previously generated as previously reported[1,2]. Spike constructs for PE.1.4, BA.3.2, and PY.1.1.1 were generated using previously reported methods[3]. All constructs were confirmed by whole-plasmid sequencing prior to pseudovirus packaging.

#### *SARS-CoV-2 pseudovirus packaging*

Pseudotyped SARS-CoV-2 was produced following a previously established protocol[4]. HEK293T cells were first transfected with spike-encoding plasmids using 1 mg/mL of PEI-MAX (Polysciences, Inc.) and cultured for 24 hours. The transfected HEK293T cells were then infected with VSV-G pseudotyped ΔG-luciferase virus (Kerafast, EH1020-PM) at a multiplicity of infection (MOI) of approximately 3 to 5. Two hours later, the cells were washed three times with and cultured in fresh competent cell medium overnight. The virus was then harvested first by centrifugation at 2000 rpm for 10 minutes followed by collection of the supernatant. 20% by volume of I1 hybridoma (ATCC, CRL-2700) supernatant was then added to each virus before storing at -80°C until further use.

*Pseudovirus neutralization*

Pseudoviruses were titrated to standardize viral input prior to each neutralization assay. For neutralization assays, serum samples were heat-inactivated at 56°C for 30 minutes before use, and the inactivated sera were serially diluted with a dilution factor of four on black Costar neutralization plates in complete culture medium. Following this, pseudoviruses were added and incubated at 37 °C for 1 hour. As a control, wells containing only the pseudovirus were also prepared on each test plate. Vero-E6 cells were then seeded at 40,000 cells per well and were incubated overnight at 37°C. Next, cell lysis was conducted and luciferase activity was quantified with Promega Luciferase Assay System (cat. E4550) and a Tecan Infinite® 200 PRO using i-control™ software v.3.9.1.0, in accordance with the manufacturer's instructions. The serum dilution that inhibits 50% of virus entry ($ID_{50}$) was calculated using a five-parameter dose-response curve fitting using R package drda v2.0.3 in R v4.3.2[5].

*Quantification and statistical analysis*

$ID_{50}$ values at or below the assay limit of detection (LOD) of 50 were treated as 50 for the purpose of calculating each group's geometric mean $ID_{50}$ titer (GMT). Half-lives were calculated using an exponential model. For individuals with no change or positive change in titer between 1- and 4-months samples, half-life of that titer was treated as 365 days for use in calculation of average half-life.

**Acknowledgements**

This study was supported by funding from the NIH SARS-CoV-2 Assessment of Viral Evolution (SAVE) Program (subcontract no. 0258-A700-4609 under federal contract no. 75N93021C00014) to D.D.H. and (subcontract GR0010139-PO024016 under federal contract no. 75N93021C00016) to A.G., the Gates Foundation (project INV019355) to D.D.H., internal startup funding UR014016 from Columbia University to Y.G., K24 AI155230 to M.T.Y., and K08 AI180347 to A.B.

We express our gratitude to Hiroshi Mohri (Columbia) for assistance with sample collection, to Jayesh G. Shah, Lawrence J. Purpura, Amanda Castillo, Meredith McNairy and Antonia Sturiza for conducting the C-PIC study (Columbia), and to Theresa Kowalski-Dobson, Anna Buswinka, Joseph Wendzinski, Mayurika Patel, Noah Paalanen and Ethan Hall of the VIVA study team for conducting the VIVA study.

**Author Contributions**

The project was conceptualized by I.A.M., Y.G., A.G., and D.D.H. Experiments were conducted and data analyzed by I.A.M., M.W., H.H., and Y.G. Serum samples were collected by I.A.M., A.B., C.G., V.M.P., M.T.Y., and A.G. The manuscript was written by I.A.M., M.W., H.H., Y.G., and D.D.H., with feedback from all authors. All contributing authors have reviewed and endorsed the manuscript.

**Declaration of Interests**

D.D.H. co-founded TaiMed Biologics and RenBio, and he serves as a consultant for WuXi Biologics and Brii Biosciences and is a board director at Vicarious Surgical. A.G. served as a member of the scientific advisory board for Janssen Pharmaceuticals and has consulted and serves on a scientific advisory board for Sanofi Pasteur. The remaining authors declare no conflicts of interest.

### Supplemental Table and Figures

| | | All participants | | Age Cohort 1 (18-49 y.o.) | | Age Cohort 2 (50-64 y.o.) | | Age Cohort 3 (65+ y.o.) | |
|---|---|---|---|---|---|---|---|---|---|
| | | No. or Mean | % or (range) | No. or Mean | % or (range) | No. or Mean | % or (range) | No. or Mean | % or (range) |
| **Total** | | 36 | - | 18 | - | 10 | - | 8 | - |
| **Female** | | 28 | 77.8% | 12 | 66.7% | 9 | 90% | 7 | 87.5% |
| **Male** | | 7 | 19.4% | 5 | 27.8% | 1 | 10% | 1 | 12.5% |
| **Prefer Not to Answer** | | 1 | 2.8% | 1 | 5.56% | - | - | - | - |
| **Age** | | 47.3 | (19, 80) | 30.4 | (19, 48) | 58.1 | (52, 63) | 71.9 | (66, 80) |
| **No. Vaccines** | All vaccines | 6.7 | (4, 10) | 5.9 | (5, 7) | 7 | (4, 8) | 8 | (4, 10) |
| | WT | 3.3 | (2, 4) | 3 | (3, 3) | 3.4 | (2, 4) | 3.6 | (2, 4) |
| | BA.5 BV | 0.8 | (0, 2) | 0.6 | (0, 1) | 1 | (0, 1) | 0.9 | (0, 2) |
| | XBB.1.5 | 0.7 | (0, 2) | 0.4 | (0, 1) | 0.8 | (0, 1) | 1.1 | (0, 2) |
| | KP. 2 MV | 0.9 | (0,2) | 0.8 | (0, 1) | 0.8 | (0, 1) | 1.4 | (1, 2) |
| | LP.8.1 MV | 1.0 | (1,1) | 1 | (1, 1) | 1 | (1, 1) | 1 | (1, 1) |
| **Sera Days Post Infection (Pre)** | | 796.0 | (0, 1626) | 874.6 | (0, 833.7) | 829.7 | (0, 1235) | 668.2 | (0, 1049) |
| **Sera Days Post Infection (Post)** | | 831.6 | (0, 1674) | 871.9 | (0, 1391) | 865.5 | (0, 1276) | 697 | (0, 1077) |
| **Sera Days Pre LP.8.1 MV Vaccination** | | 6.1 | (0, 28) | 6.6 | (0, 28) | 6.8 | (1, 16) | 4.1 | (0, 25) |
| **Sera Days Post LP.8.1 MV Vaccination** | | 28.6 | (21, 45) | 30.6 | (21, 45) | 26.3 | (22, 34) | 27 | (23, 33) |

**Table S1. Summary of clinical cohort for pre- and post-boost analysis.**
No., number; y.o., years old; WT, wildtype; MV, monovalent vaccine; BV, bivalent vaccine.

| ID | Age (Yr) | Sex | Race | No. Vax | No. WT Vax | No. BA.5 Bivalent Vax | No XBB.1.5 Vax | No. KP.2 MV | No. LP.8.1 MV | Sera Days Post Most Recent Infx Pre | Sera Days Post Most Recent Infx Post | Sera Days Pre LP.8.1 MV | Sera Days Post LP.8.1 MV | Vaccine History |
|---|---|---|---|---|---|---|---|---|---|---|---|---|---|---|
| CUMC 1 | 26 | F | Asian | 7 | 3 | 1 | 1 | 1 | 1 | NA | NA | 2 | 28 | WT-P/WT-P/WT-P/BA.5-M/XBB.1.5-M/KP.2-M/LP.8.1-M |
| CUMC 3 | 22 | M | Asian | 5 | 3 | 0 | 0 | 1 | 1 | 141 | 175 | 3 | 31 | WT-P/WT-P/WT-P/KP.2-P/LP.8.1-P |
| CUMC 8 | 23 | F | Asian | 5 | 3 | 0 | 0 | 1 | 1 | NA | NA | 1 | 31 | WT-P/WT-P/WT-P/KP.2-M/LP.8.1-M |
| CUMC 16 | 23 | M | Asian | 4 | 3 | 0 | 0 | 0 | 1 | 1030 | 1063 | 1 | 32 | WT-P/WT-P/WT-P/LP.8.1-P |
| CUMC 18 | 23 | F | Asian | 4 | 3 | 0 | 0 | 0 | 1 | NA | NA | 2 | 31 | WT-S/WT-S/WT-S/LP.8.1-P |
| MICH 1 | 48 | F | White | 5 | 3 | 0 | 0 | 0 | 1 | 896 | 927 | 0 | 31 | WT-P/WT-P/WT-P/XBB.1.5-P/LP.8.1-P |
| MICH 2 | 30 | F | Black or African American | 6 | 3 | 1 | 0 | 1 | 1 | 1107 | 1148 | 7 | 34 | WT-P/WT-P/WT-P/BA.5-P/KP.2-P/LP.8.1-P |
| MICH 3 | 31 | M | White | 6 | 3 | 0 | 1 | 1 | 1 | 901 | 953 | 28 | 24 | WT-P/WT-P/WT-P/XBB.1.5-P/KP.2-P/LP.8.1-M |
| MICH 4 | 75 | F | More than One Race | 10 | 4 | 2 | 1 | 2 | 1 | NA | NA | 1 | 26 | WT-M/WT-M/WT-M/WT-M/BA.5-M/BA.5-M/XBB.1.5-M/KP.2-N/KP.2-P/LP.8.1-M |
| MICH 5 | 33 | F | White | 7 | 3 | 1 | 1 | 1 | 1 | 260 | 286 | 1 | 25 | WT-P/WT-P/WT-P/BA.5-P/XBB.1.5-P/KP.2-M/LP.8.1-M |
| MICH 6 | 26 | F | White | 5 | 3 | 1 | 0 | 0 | 1 | 1072 | 1115 | 9 | 34 | WT-P/WT-P/WT-P/BA.5-P/LP.8.1-M |
| MICH 7 | 52 | F | White | 7 | 3 | 1 | 1 | 1 | 1 | NA | NA | 10 | 25 | WT-P/WT-P/WT-P/BA.5-P/XBB.1.5-M/KP.2-M/LP.8.1-M |
| MICH 8 | 55 | F | White | 8 | 4 | 1 | 1 | 1 | 1 | 742 | 778 | 4 | 32 | WT-P/WT-P/WT-P/WT-P/BA.5-P/XBB.1.5-P/KP.2-N/LP.8.1-M |
| MICH 9 | 29 | F | White | 6 | 3 | 1 | 0 | 1 | 1 | 430 | 473 | 9 | 34 | WT-P/WT-P/WT-P/BA.5-U/KP.2-U/LP.8.1-P |
| MICH 10 | 60 | F | White | 7 | 3 | 1 | 1 | 1 | 1 | 1235 | 1276 | 7 | 34 | WT-M/WT-M/WT-M/BA.5-M/XBB.1.5-M/KP.2-M/LP.8.1-M |
| MICH 11 | 60 | F | White | 6 | 3 | 1 | 0 | 1 | 1 | 1177 | 1220 | 16 | 27 | WT-M/WT-M/WT-M/BA.5-P/KP.2-M/LP.8.1-M |
| MICH 12 | 29 | F | White | 7 | 3 | 1 | 1 | 1 | 1 | 925 | 964 | 8 | 31 | WT-P/WT-P/WT-P/BA.5-P/XBB.1.5-P/KP.2-P/LP.8.1-P |
| MICH 13 | 38 | F | White | 7 | 3 | 1 | 1 | 1 | 1 | 824 | 856 | 0 | 32 | WT-P/WT-P/WT-P/BA.5-P/XBB.1.5-P/KP.2-P/LP.8.1-M |
| MICH 14 | 63 | F | White | 4 | 2 | 1 | 0 | 0 | 1 | NA | NA | 1 | 22 | WT-P/Wt-M/BA.5-M/LP.8.1-P |
| MICH 15 | 46 | F | White | 6 | 3 | 1 | 0 | 1 | 1 | 1626 | 1674 | 3 | 45 | WT-P/WT-P/WT-P/BA.5-P/KP.2-P/LP.8.1-P |
| MICH 16 | 37 | F | White | 7 | 3 | 1 | 1 | 1 | 1 | 190 | 239 | 28 | 21 | WT-P/WT-P/WT-P/BA.5-P/XBB.1.5-P/KP.2-P/LP.8.1-P |
| MICH 17 | 59 | M | White | 8 | 4 | 1 | 1 | 1 | 1 | NA | NA | 3 | 24 | WT-M/WT-M/WT-M/WT-M/BA.5-M/XBB.1.5-M/KP.2-M/LP.8.1-M |
| MICH 18 | 72 | M | White | 10 | 4 | 1 | 2 | 2 | 1 | 479 | 502 | 0 | 23 | WT-M/WT-M/WT-M/WT-M/BA.5-M/XBB.1.5-M/XBB.1.5-N/KP.2-M/KP.2-M/LP.8.1-M |
| MICH 19 | 33 | O | White | 7 | 3 | 1 | 1 | 1 | 1 | 748 | 784 | 2 | 34 | WT-M/Wt-M/WT-M/BA.5-U/XBB.1.5-U/KP.2-U/LP.8.1-M |
| MICH 20 | 57 | F | White | 6 | 3 | 1 | 1 | 0 | 1 | 1110 | 1138 | 1 | 27 | WT-P/WT-P/WT-M/BA.5-M/XBB.1.5-P/LP.8.1-M |
| MICH 21 | 31 | M | White | 5 | 3 | 0 | 0 | 1 | 1 | 1355 | 1391 | 13 | 23 | WT-P/WT-P/WT-M/KP.2-M/LP.8.1-P |
| MICH 22 | 19 | M | More than One Race | 7 | 3 | 1 | 1 | 1 | 1 | 1000 | 1031 | 1 | 30 | WT-P/WT-P/WT-P/BA.5-P/XBB.1.5-P/KP.2-M/LP.8.1-P |
| MICH 23 | 73 | F | White | 8 | 4 | 1 | 1 | 1 | 1 | 197 | 228 | 1 | 30 | WT-P/WT-P/WT-P/WT-P/BA.5-P/XBB.1.5-M/KP.2-M/LP.8.1-M |
| MICH 24 | 66 | F | White | 10 | 4 | 1 | 2 | 2 | 1 | 576 | 609 | 0 | 33 | WT-P/WT-P/WT-P/WT-P/BA.5-P/XBB.1.5-P/XBB.1.5-P/KP.2-M/KP.2-P/LP.8.1-M |
| MICH 25 | 60 | F | White | 8 | 4 | 1 | 1 | 1 | 1 | 480 | 510 | 7 | 23 | WT-M/WT-M/WT-M/WT-M/BA.5-M/XBB.1.5-P/KP.2-M/LP.8.1-P |
| MICH 26 | 73 | F | White | 7 | 3 | 1 | 1 | 1 | 1 | N/A | N/A | 25 | 24 | WT-P/WT-P/WT-P/BA.5-P/XBB.1.5-P/KP.2-P/LP.8.1-P |
| MICH 27 | 80 | F | White | 8 | 4 | 1 | 1 | 1 | 1 | 1049 | 1077 | 1 | 27 | WT-P/WT-P/WT-P/WT-P/BA.5-M/XBB.1.5-M/KP.2-M/LP.8.1-M |
| MICH 28 | 56 | F | Asian | 8 | 4 | 1 | 1 | 1 | 1 | NA | NA | 4 | 27 | WT-M/WT-M/WT-M/WT-M/BA.5-P/XBB.1.5-M/KP.2-N/LP.8.1-M |
| MICH 29 | 68 | F | White | 7 | 4 | 0 | 1 | 1 | 1 | 666 | 696 | 0 | 30 | WT-P/WT-P/WT-M/WT-M/XBB.1.5-M/KP.2-M/LP.8.1-M |
| MICH 30 | 68 | F | White | 4 | 2 | 0 | 0 | 1 | 1 | 1042 | 1070 | 5 | 23 | WT-J/WT-M/KP.2-M/LP.8.1-M |
| MICH 31 | 59 | F | White | 8 | 4 | 1 | 1 | 1 | 1 | 234 | 271 | 15 | 22 | WT-M/WT-M/WT-M/WT-M/BA.5-M/XBB.1.5-P/KP.2-P/LP.8.1-M |

**Table S2. Demographics of clinical cohort for pre- and post-boost analysis.**

Vaccine formulations are denoted as wildtype (WT), BA.5 Bivalent (BA.5), XBB.1.5 monovalent (XBB.1.5), and KP.2 monovalent (KP.2). Vaccine manufacturers are denoted as Pfizer (P), Moderna (M), and Unknown (U). Yr, years; Infx, infection; Vax, vaccination; DBV, days before vaccination; DPV, days post vaccination; DPI, days post infection; F, female; M, male; Wh, white; Af, Black or African American; As, Asian.

| | | All participants | |
|---|---|---|---|
| | | No. or Mean | % or (range) |
| **Total** | | 11 | - |
| **Female** | | 6 | 54.5% |
| **Male** | | 4 | 36.4% |
| **Prefer Not to Answer** | | 1 | 9.1% |
| **Age** | | 36.0 | (22, 72) |
| **No. Vaccines** | All vaccines | 6.3 | (4, 10) |
| | WT | 3.2 | (2, 4) |
| | BA.5 BV | 0.6 | (0, 2) |
| | XBB.1.5 | 0.6 | (0, 2) |
| | KP. 2 MV | 0.8 | (0,2) |
| | LP.8.1 MV | 1.0 | (1,1) |
| **Sera Days Post Infection (1m)** | | 794.3 | (0, 1291) |
| **Sera Days Post Infection (4m)** | | 869.9 | (0, 1354) |
| **Sera Days Pre LP.8.1 MV Vaccination (1m)** | | 29.5 | (24, 36) |
| **Sera Days Post LP.8.1 MV Vaccination (4m)** | | 111.7 | (85, 156) |

**Table S3: Summary of clinical cohort for durability post-boost analysis.**

No., number; y.o., years old; WT, wildtype; MV, monovalent vaccine; BV, bivalent vaccine; 1m, ~1 month post-boost sample; 4m, ~4 month post-boost sample.

| ID | Age (Yr) | Sex | Race | No. Vax | No. WT Vax | No. BA.5 Bivalent Vax | No XBB.1.5 Vax | No. KP.2 MV | No. LP.8.1 MV | Sera Days Post Most Recent Infx (1m) | Sera Days Post Most Recent Infx (4m) | Sera Days Post LP.8.1 MV (1m) | Sera Days Post LP.8.1 MV (4m) | Vaccine History |
|---|---|---|---|---|---|---|---|---|---|---|---|---|---|---|
| CUMC 1 | 26 | F | Asian | 7 | 3 | 1 | 1 | 1 | 1 | NA | NA | 28 | 152 | WT-P/WT-P/WT-P/BA.5-M/XBB.1.5-M/KP.2-M/LP.8.1-M |
| CUMC 3 | 22 | M | Asian | 5 | 3 | 0 | 0 | 1 | 1 | 175 | 300 | 31 | 156 | WT-P/WT-P/WT-P/KP.2-P/LP.8.1-P |
| CUMC 8 | 23 | F | Asian | 5 | 3 | 0 | 0 | 1 | 1 | NA | NA | 31 | 142 | WT-P/WT-P/WT-P/KP.2-M/LP.8.1-M |
| CUMC 16 | 23 | M | Asian | 4 | 3 | 0 | 0 | 0 | 1 | 1063 | 1173 | 32 | 142 | WT-P/WT-P/WT-P/LP.8.1-P |
| CUMC 17 | 22 | F | Asian | 5 | 3 | 0 | 0 | 1 | 1 | 1291 | 1354 | 36 | 99 | WT-P/WT-P/WT-P/KP.2-P |
| MICH 5 | 33 | F | White | 7 | 3 | 1 | 1 | 1 | 1 | 286 | 357 | 25 | 96 | WT-P/WT-P/WT-P/BA.5-P/XBB.1.5-P/KP.2-M/LP.8.1-M |
| MICH 6 | 26 | F | White | 5 | 3 | 1 | 0 | 0 | 1 | 1115 | 1171 | 34 | 90 | WT-P/WT-P/WT-P/BA.5-P/LP.8.1-M |
| MICH 17 | 59 | M | White | 8 | 4 | 1 | 1 | 1 | 1 | NA | NA | 24 | 88 | WT-M/WT-M/WT-M/WT-M/BA.5-M/XBB.1.5-M/KP.2-M/LP.8.1-M |
| MICH 18 | 72 | M | White | 10 | 4 | 1 | 2 | 2 | 1 | 502 | 568 | 23 | 89 | WT-M/WT-M/WT-M/WT-M/BA.5-M/XBB.1.5-M/XBB.1.5-N/KP.2-M/KP.2-M/LP.8.1-M |
| MICH 19 | 33 | O | White | 7 | 3 | 1 | 1 | 1 | 1 | 784 | 840 | 34 | 90 | WT-M/Wt-M/WT-M/BA.5-U/XBB.1.5-U/KP.2-U/LP.8.1-M |
| MICH 20 | 57 | F | White | 6 | 3 | 1 | 1 | 0 | 1 | 1138 | 1196 | 27 | 85 | WT-P/WT-P/WT-M/BA.5-M/XBB.1.5-P/LP.8.1-M |

**Table S4. Demographics of clinical cohort for durability post-boost analysis.**

Vaccine formulations are denoted as wildtype (WT), BA.5 Bivalent (BA.5), XBB.1.5 monovalent (XBB.1.5), and KP.2 monovalent (KP.2). Vaccine manufacturers are denoted as Pfizer (P), Moderna (M), and Unknown (U). Yr, years; Infx, infection; Vax, vaccination; DBV, days before vaccination; DPV, days post vaccination; DPI, days post infection; F, female; M, male; Wh, white; Af, Black or African American; As, Asian.

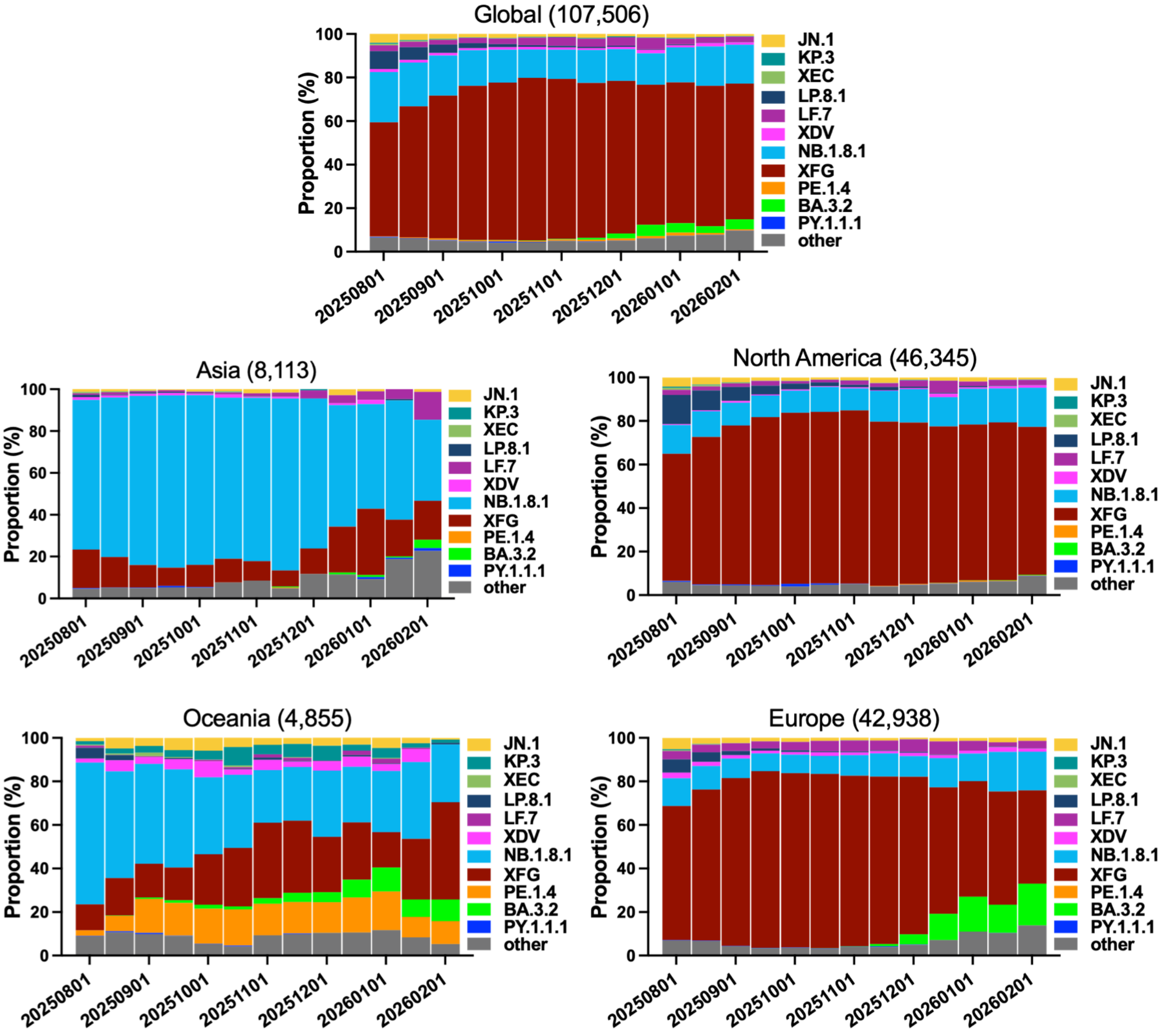

**Figure S1. Frequencies of SARS-CoV-2 variants around the world.**

Relative frequencies of SARS-CoV-2 variants from 08/01/2025 to 02/01/2026 in the indicated regions. Numbers of sequences analyzed in parentheses above each subpanel. Data from GISAID.

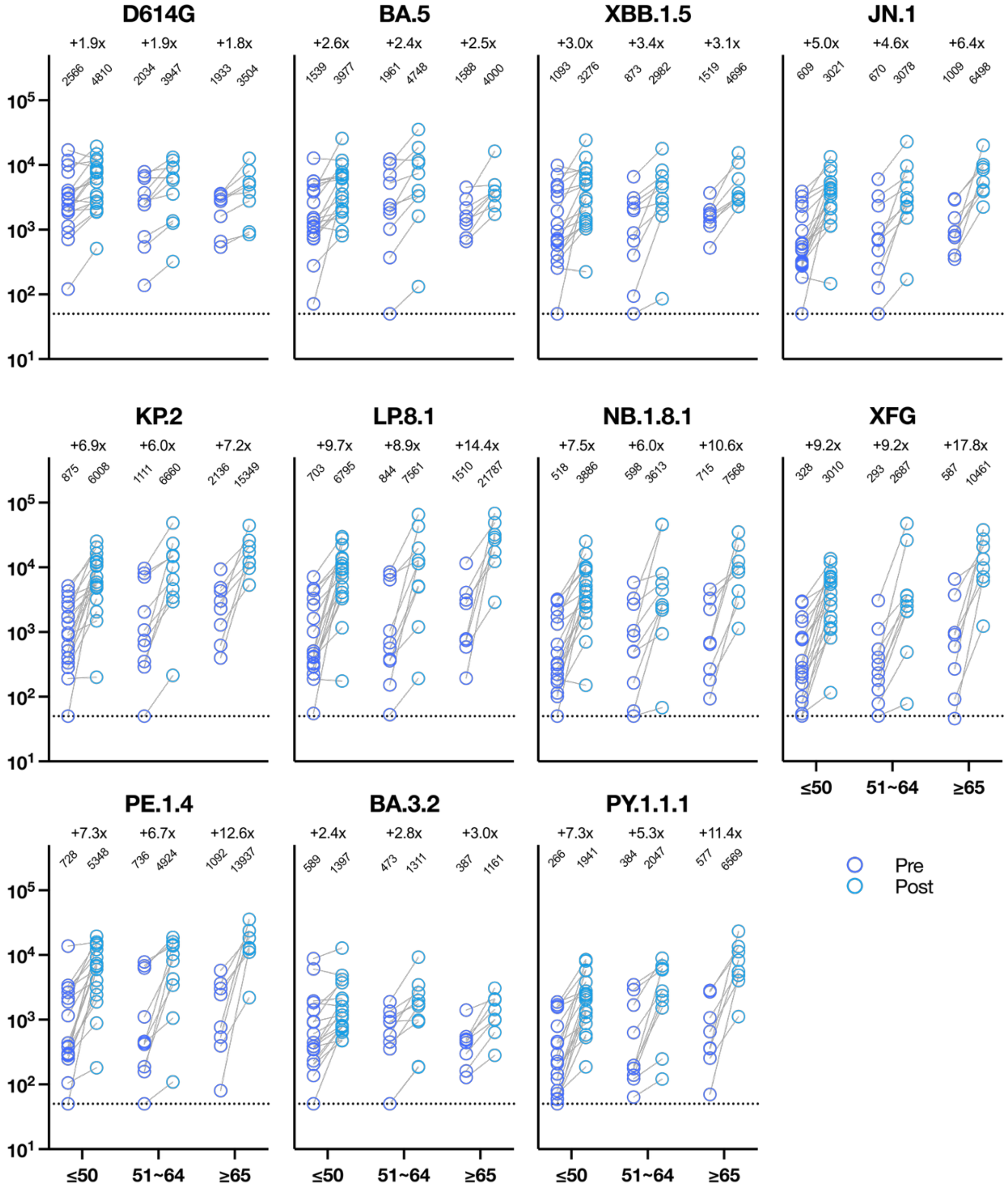

**Figure S2. LP.8.1 MV boost effects by KP.2 MV between age groups.**

Data are presented as fold changes in neutralization $ID_{50}$ titers following LP.8.1 MV vaccination for across age groups. Geometric mean titers (GMT) are shown above each sample set, and the fold change from pre- to post-booster is shown above GMTs. MV, monovalent vaccine. n, sample size. The dotted line represents the assay limit of detection (LOD) of 50.
